# A novel insertional allele of *CG18135* gene is associated with severe mutant phenotypes in *Drosophila melanogaster*

**DOI:** 10.1101/2022.09.13.507739

**Authors:** Attila Cristian Ratiu, Adrian Ionascu, Alexandru Al. Ecovoiu

**Affiliations:** Drosophila Laboratory, Faculty of Biology, University of Bucharest, Intrarea Portocalelor, No. 1-3, 060101, Bucharest, Romania; Academy of Romanian Scientists, Ilfov, No. 3, 050044, Bucharest, Romania

**Keywords:** CG18135, P{lacW} insertion, Drosophila melanogaster, mutant phenotype, lethality, gene expression

## Abstract

*Drosophila melanogaster* has been at the forefront of genetic studies and biochemical modelling for over a century. Yet, the functions of many genes are still unknown mainly because no phenotypic data are available. Herein, we present first evidence data regarding the particular molecular and other quantifiable phenotypes, such as viability and anatomical anomalies, induced by a novel *P{lacW}* insertional mutant allele of *CG18135* gene. So far, the *CG18135* functions have only been theorized based on electronic annotation and presumptive associations inferred upon high-throughput proteomics or RNA sequencing experiments. The descendants of individuals harboring the *CG18135*^*P{lacW}CG18135*^ allele were scored in order to assess mutant embryos, larvae and pupae viability versus Canton Special. Our results revealed that the homozygous *CG18135*^*P{lacW}CG18135*^/*CG18135*^*P{lacW}CG18135*^ genotype determines significant lethality both at the inception of larval stage and during pupal development. Few imago escapers that breach the puparium and even more rarely fully exit from it exhibit specific eye depigmentation, wing abnormal unfolding and strong locomotor impairment with apparent spasmodic legs movements. Their maximum lifespan is shorter than two days. When using the quantitative Real-Time PCR (qRT-PCR) method to confirm that *CG18135* is indeed upregulated in males compared to females an unexpected gene upregulation was also detected in heterozygous mutants comparative to wild-type flies, probably because of regulatory perturbations induced by *P{lacW}* transposon. Our work provides the first phenotypic evidence for the essential role of *CG18135*, a scenario in accordance with the putative role of this gene in the carbohydrate binding processes.

## Introduction

*Drosophila melanogaster* has been studied for over a century, being the definitive model organism in genetics, biochemistry, molecular biology and bioinformatics. Even if the *D. melanogaster* genome was sequenced in 2000 (Adams *et al*., 2000), for a number of computed genes (CG) there are no available phenotypic data in support of their characterization and annotation. One example is *CG18135* which was identified in 2004 via computational techniques (Gramates *et al*., 2022).

*CG18135* has the gene sequence location 3L:18,990,026..18,998,283, is located on the genomic minus strand and is known to encode for five transcripts which are translated in proteins having between 636 and 770 amino acids. *CG18135* sequence is conserved in higher eukaryotes such as fish, reptiles, birds, and mammals, but also in invertebrates and yeasts, according to DIOPT v9.1 (Hu *et al*., 2011). *CG18135* gene is orthologous to human *glycerophosphocholine phosphodiesterase 1* (*GPCPD1*). Whilst the function of the *CG18135* gene was not previously genetically studied in *D. melanogaster*, the associated protein was evidenced to facilitate myosin filament binding during embryogenesis via interaction with the Myo10A protein at the plasma membrane level (Liu *et al*., 2008) and to participate in glycerophospholipids catabolism (Gaudet *et al*., 2011).

*CG18135* has been documented to participate in embryogenesis and to be active through all the developmental stages in *D. melanogaster*. Increased levels of CG18135 RNA were identified in both early (0-4 hours) and late (20-24 hours) embryos (Fisher *et al*., 2012). Regarding other developmental stages, highly increased levels of CG18135 RNA were found in pupae and in adult flies compared to embryos, suggesting that the gene plays an important function beyond the embryo stage (Casas-Vila *et al*., 2017). Moreover, it has been shown that one week old adult males have significantly higher *CG18135* gene expression than similarly aged females (Brown *et al*., 2014; Casas-Vila *et al*., 2017).

Several transposon insertions in *CG18135* were previously reported in FlyBase (FB2023_05), namely nine *P*, one *PiggyBac* (*PBac*) and two *Minos* (*MiMIC*) derived artificial mobile elements (Gramates *et al*., 2022; Buels *et al*., 2016). The *P* and *PBac* artificial transposons, as well as one *MiMIC* (Venken *et al*., 2011) are located in the 5’ half of *CG18135*, defining a local insertional hotspot for 11 transposons in a 1267 bp span. Curiously, none of them determine noticeable phenotypic consequences.

During one of our previously experiments we achieved mutagenesis mediated by the mobilization of *P{lacW}γCop*^*057302*^ transposon (Ratiu *et al*., 2008) located in 5’UTR of *γCop*. The idea backing this study was that transposon mobilization starting from an essential gene, such as *γCop*, will likely lead to conservative reinsertions in other essential genes, eventually functionally linked. Thus, we generated the MZ4CM3 line with individuals that contain both *P{lacW}γCop*^*057302*^ and *P{lacW}*^*CG18135*.*MZ4CM3*^ transposons located in the 3R and, respectively, 3L chromosomes. The latter *P{lacW}* resides in the first intron of CG18135-RB transcript at the genomic coordinates 3L:18996755..18996762 (GenBank/NCBI accession number: HQ695001.1). The aforementioned genomic interval indicates the localization of the characteristic duplicated octet, since this is actually the maximum resolution for P transposon insertions mapping, as debated elsewhere (Ecovoiu *et al*., 2016). Since P{lacW}γCop^057302^ determines embryo lethality in homozygous condition (Deak *et al*., 1997; Ecovoiu *et al*., 2002), the actual phenotypic effect of *P{lacW}*^*CG18135*.*MZ4CM3*^ was impossible to assess as long as the two alleles were linked. Only recently we successfully performed a series of genetic crosses allowing us to separate the *P{lacW}γCop*^*057302*^ and *P{lacW}*^*CG18135*.*MZ4CM3*^ transposons by crossing-over and to construct *CG18135*^*P{lacW}CG18135*^-Sep1H line uniquely containing the novel *P{lacW}* insertion in *CG18135*.

Herein we report the first phenotypic and functional characterization of this original insertional allele of *CG18135* gene, symbolized *CG18135*^*P{lacW}CG18135*^. Our main objectives were to investigate the viability of heterozygous and homozygous *CG18135*^*P{lacW}CG18135*^ mutant flies during various developmental stages, to score the phenotypic features of the elusive homozygous mutant adult escapers, to evaluate the *CG18135* expression level in adult male and female mutant heterozygotes. Moreover, we used a set of bioinformatics tools in order to investigate the putative regulatory consequences induced by the *P{lacW}* alien.

## Materials and methods

### D. melanogaster lines and rearing

In this study, two distinct *D. melanogaster* lines were used: *CG18135*^*P{lacW}CG18135*^ mutant line and the reference Canton Special (CantonS) line. The mutant flies harbor at least one insertional allele caused by the *P{lacW}* insertion in the 5’ region of the *CG18135* gene (GenBank/NCBI accession number: HQ695001.1). The mutant allele was kept over two distinct balancer chromosomes and the respective derived lines were symbolized as it follows:

- *y*^-^, *w*^-^; *CG18135*^*P{lacW}CG18135*^*-Sep1H* / TM3, *Ser, GFP, e*; in brief, *CG18135*^*P{lacW}*^/GFP;
- *w*^*-*^; *CG18135*^*P{lacW}CG18135*^*-Sep1H* / TM6B, *Tb, Hu, e*; in brief, *CG18135*^*P{lacW}*^/TM6B.

The flies were placed in glass vials on standard wheat-banana-yeast-agar based growth medium and kept under 12h/12h light/dark cycles, constant temperature of 18°-19°C and humidity of 20-60%. Under experimental conditions, the flies were raised in plastic vials or Petri dishes on the standard growth medium and alternative on the commercially available Grape Agar medium (Genesee Scientific).

### Viability experiment

Flies from the *CG18135*^*P{lacW}*^/GFP line and CantonS line were placed for a maximum of 24 hours in 50 mm embryo collection cages (FlyStuff) coupled with Petri dishes containing Grape Agar medium (roughly 100 males and 100 females/cage). Throughout the developmental stages, the fluorescent phenotype, determined by the presence or absence of the *GFP* construct, was assessed by using an Olympus U-RFL-T UV lamp. The collected embryos were placed individually or in groups of five in 24 well-plates on standard medium. Firstly, the focus was on monitoring the embryo development and for this purpose we used the former experimental design and each well was regularly inspected and counted until second or third instar larvae were noticed. For this particular evaluation, the *CG18135*^*P{lacW}*^/TM6B line was also used, following the same procedures. A second round of experiments was mainly focused on mutant homozygous individuals; the surviving non-GFP larvae from each group of five were removed and individually placed in 48 well-plates containing standard medium. Here, they were inspected at least two times a day during their larval and pupal development. Rare adult individuals that emerged from puparium were phenotypically scored and pictures as well as short videos were taken.

### Gene expression experiment

*CG18135* gene expression was measured in 50-hour old adult females and males from *CG18135*^*P{lacW}*^/TM6B mutant line and CantonS control line. Biological triplicates consisting of 20 flies from each sex were treated for RNA stabilization using the DNA/RNA Shield kit, then submitted to RNA extraction using the RNeasy Mini Kit (Promega). cDNA conversion was performed combining the SuperScript II Reverse Transcriptase (Invitrogen) with the Reverse Transcription System kit (Promega) using random primers and adjusting the RNA volumes based on RNA concentrations.

qRT-PCR amplifications were performed in technical triplicates using the SYBR Green ROX qPCR Mastermix kit (Qiagen) with newly designed *CG18135* primers (Table 1) and literature available *RpL32* endogene primers (Fiumera *et al*., 2005). Each qRT-PCR reaction tube contained 20 ng of cDNA, 0.5 µM of primers (for each forward/reverse primer), 1X Mastermix solution and PCR-grade water up to 20 µL final reaction volume. The cycle threshold (Ct) values were gathered at a fluorescence threshold Rn = 0.2.

**Table 1.**
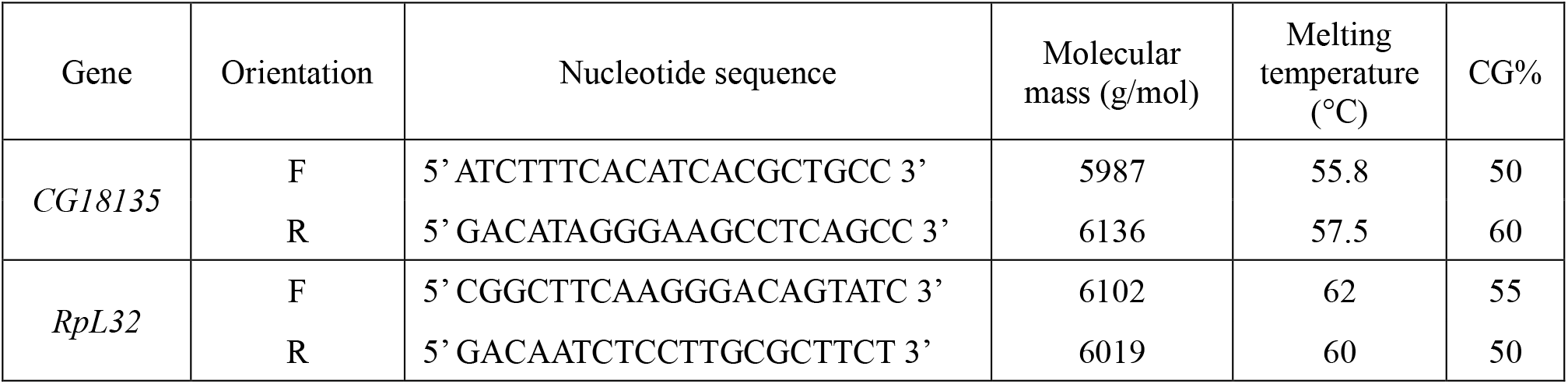
Characteristics of the primers used for gene expression analysis with qRT-PCR. The primers’ melting temperatures were provided by Integrated DNA Technologies for *CG18135* and Eurogentec for *RpL32*.

### Data analysis

Microsoft Excel was used for data manipulation of raw Ct values collected from the qRT-PCR experiment. Statistical analysis of data collected during the viability experiments was performed using either Microsoft Excel, RStudio Desktop (version ≥ 4.2.0) or GraphPad Prism 8.0.2 for Windows (GraphPad Software, Boston, Massachusetts USA, www.graphpad.com). For the gene expression data analysis, we implemented the Livak formula (Livak and Schmittgen, 2001). Statistical testing on 2^-ΔCt^ values and relative expression fold change (FC) were evaluated using the qDATA R application (https://github.com/A-Ionascu/qDATA). Graphical representations were created with GraphPad Prism. Statistical significance was considered at a p value ≤ 0.05.

The genomic context of *CG18135* was visualized with JBrowse 1.16.10 accessed from FlyBase and using the BLAST, Nucleotide View, Gene span, RNA, Transgenic Insertions, and Transcriptional Regulatory Regions (REDfly) tracks, as well as Set highlight from View options. The cis-regulatory modules (CRMs) are estimated according to REDfly v9.6.2 (Rivera *et al*., 2018). Motif identification within the considered CRMs was performed with the XSTREME tool from the MEME Suite 5.5.4 (Grant and Bailey, 2021). For subsequent inquiries regarding the known functions of the proteins that bind specific motifs we used FlyBase data.

## Results

### Viability experiment

In the first viability experiment we collected a total of 117, 176 and 236 embryos corresponding to CantonS, *CG18135*^*P{lacW}*^/TM6B and *CG18135*^*P{lacW}*^/GFP, respectively. After we randomly placed individual embryos in the 24 available wells of each plate, we appraised their development until they reached the second or third instar larval stages. We recorded embryonic and larval lethality within each *D. melanogaster* line and assessed the fluorescent phenotype of viable larvae from the *CG18135*^*P{lacW}*^/GFP line (Table 2). For embryos, we estimated the survival rate by subtracting the number of observed dead embryos from their initial total. Regarding the larval lethality, it was mainly assessed on the indirect evidences found in each corresponding well such as empty vitelline coat, moth pieces residuals, foraging evidences or, in some wells, their absence.

**Table 2.** In this table there are the data collected in the first experiment and the number of dead embryos, observed larvae and presumably dead larvae are indicated.

| Development | Phenotype | CantonS | *CG18135<sup>P{lacW}</sup>/TM6B* | *CG18135<sup>P{lacW}</sup>/GFP* |
| --- | --- | --- | --- | --- |
| Total embryos |  | 117 | 176 | 236 |
| Embryonic lethality | non-GFP | 7 | 69 | 54 |
|  | GFP | - | - | 33 |
| Larval viability | non-GFP | 106 | 72 | 40 |
|  | GFP | - | - | 71 |
| Larval lethality |  | 4 | 35 | 38 |
| Total larvae |  | 110 | 107 | 149 |

For the CantonS embryos we observed a mortality rate of ≈ 6%, a figure close to an alternative embryonic lethality calculation of 8.8%; the former value was considered as a baseline embryonic lethality and was used for normalizing lethality data collected for the mutant lines. The overall mutant embryo lethality was ≈ 37% for *CG18135*^*P{lacW}*^/GFP and ≈ 39% for *CG18135*^*P{lacW}*^/TM6B, considerably higher than that recorded for the control, but care should be taken when interpreting these results. The maneuvered embryos were the product of mattings between *CG18135*^*P{lacW}*^/GFP (or *CG18135*^*P{lacW}*^/TM6B) individuals, therefore there were three theoretically expected descendant genotypes, *CG18135*^*P{lacW}*^/GFP (50%), *CG18135*^*P{lacW}*^*/CG18135*^*P{lacW}*^ (25%) and GFP/GFP (25%). The latter category exhibits complete lethality during early embryonic stages; therefore, we anticipate a total embryo lethality of roughly 25% to which we have to add 4.5% (a correction relative to CantonS data). The same approach is applicable to TM6B/TM6B embryos.

In practice, statistical analysis (Figure 1A) revealed no significant difference between the observed (87 embryos) and the expected (70 embryos) embryonic lethality estimated for the descendants of the mutant *CG18135*^*P{lacW}*^/GFP individuals (χ^2^ = 1.32, 1, and p = 0.2397 for the chi-square test, and p = 0.2688 for the Fisher’s exact test). It is plausible that among the dead embryos there were some pertaining to *CG18135*^*P{lacW}*^/GFP and/or *CG18135*^*P{lacW}*^*/CG18135*^*P{lacW}*^ genotypes, but we expect that the GFP/GFP genotype hold up the majority of embryonic lethality.

**Figure 1.**
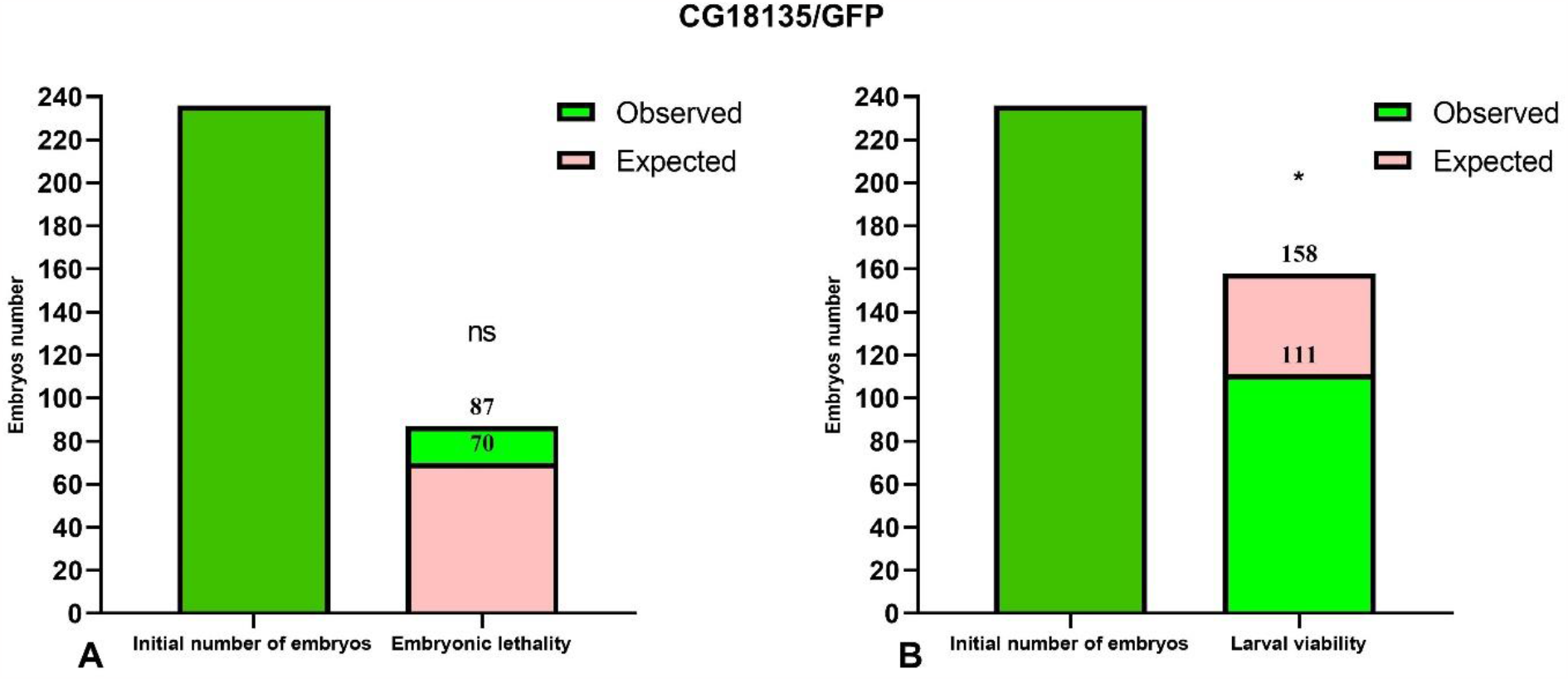
Barplots of relative embryonic lethality (A) and larval viability (B) relative to the initial number of embryos considered for *CG18135*^*P{lacW}*^/GFP line. The observed total lethality of embryos (87 embryos) is compared to the expected 29.5% lethality (70 embryos). Likewise, the observed larval viability (111) is opposed to the expected viability (158). Both chi-square and Fisher’s exact tests failed to find statistical significance (ns, because calculated p values are > 0.05) for embryonic lethality, but calculated statistically significant differences for larval viability (* for 0.01 < p < 0.05).

Regarding the observed embryonic lethality evaluated for *CG18135*^*P{lacW}*^/TM6B individuals, we also did not find statistical significance (χ^2^ = 1.780, 1, and p = 0.1822 for the chi-square test, and p = 0.2059 for the Fisher’s exact test).

Considering the embryonic lethality, we assumed that all the viable embryos continued their development throughout larval stages. During the first viability evaluation we found a second and third instar larvae viability of ≈ 41%, 47% and 90.6% (from the initial total embryos) for *CG18135*^*P{lacW}*^/TM6B, *CG18135*^*P{lacW}*^/GFP and CantonS, respectively. These percentages were mirrored by larval lethality data of ≈ 19.9%, 16.1% and 3.4% collected for the same aforementioned lines. If we take into account the CantonS larval lethality, then the expected larval viability for the mutant lines is approximatively 70.5 - 3.4 = 67.1%. In this context, there are statistically significant differences between expected and observed larval viability for both *CG18135*^*P{lacW}*^/GFP (χ^2^ = 5.252, 1, and p = 0.0219 for the chi-square test, and p = 0.0263 for the Fisher’s exact test; Figure 1B) and *CG18135*^*P{lacW}*^/TM6B (χ^2^ = 7.285, 1, and p = 0.007 for the chi-square test, and p = 0.0087 for the Fisher’s exact test) mutant lines. The lowered viability for *CG18135*^*P{lacW}*^/GFP is not significantly different from that of *CG18135*^*P{lacW}*^/TM6B (χ^2^ = 0.5272, 1 and p = 0.4678 for the chi-square test, and p = 0.4988 for the Fisher’s exact test). Moreover, neither larval nor pupal mortality was significantly different when comparing *CG18135*^*P{lacW}*^/GFP to CantonS offspring.

For the second viability evaluation we focused on *CG18135*^*P{lacW}*^/GFP line and started form a total number of 765 embryos. Batches of five randomly selected embryos were placed in each of the 24 wells of a given culture plate. When larval viability was estimated, we assumed by default an embryonic lethality of 37%, as previously estimated. In accordance, we counted a viability of ≈ 43.8%, which is very close to our prior numbers. Out of the viable second and third instar larvae, we selected 104 non-GFP larvae that were individually placed in 48-wells plates. Each well was monitored on a daily basis and the developmental progressions were carefully noted. We found that ≈ 14.4% of the larvae do not reach the pupal stage, while all the remaining ones proceed at least until late pupal stages. Compared with the initial number of larvae, around 65% of them get fully developed eyes and folded wings visible through the puparium. Out of the starting larvae, at least 12.5% attain the imago stage and the corresponding adults either only open the puparium (Figure 1 A and B, Supplementary file 1) and are eventually partially getting out of it, or completely breach the puparium and land on medium. Either way, the maximum lifespan of the partially or fully emerged imago is about 48 hours.

### The imago mutant phenotype

The escaper imago individuals of both sexes that harbor the *CG18135*^*P{lacW}CG18135*^/*CG18135*^*P{lacW}CG18135*^ genotype exhibit a series of noteworthy specific phenotypes beside the early onset lethality. The firstly noted is a peculiar central depigmentation visible at eyes level. Our individuals have *w*^*-*^ genetic background, therefore their eye color is caused by the *mini-white*^*+*^ allele present in the *P{lacW}* transposon. This depigmentation is apparent in both puparium captive and emerged adults (Figure 2). When naturally or manually puparium-free imago individuals were available, we also noticed an abnormal wing folding during the maximum life span of two days (Figure 2C).

**Figure 2.**
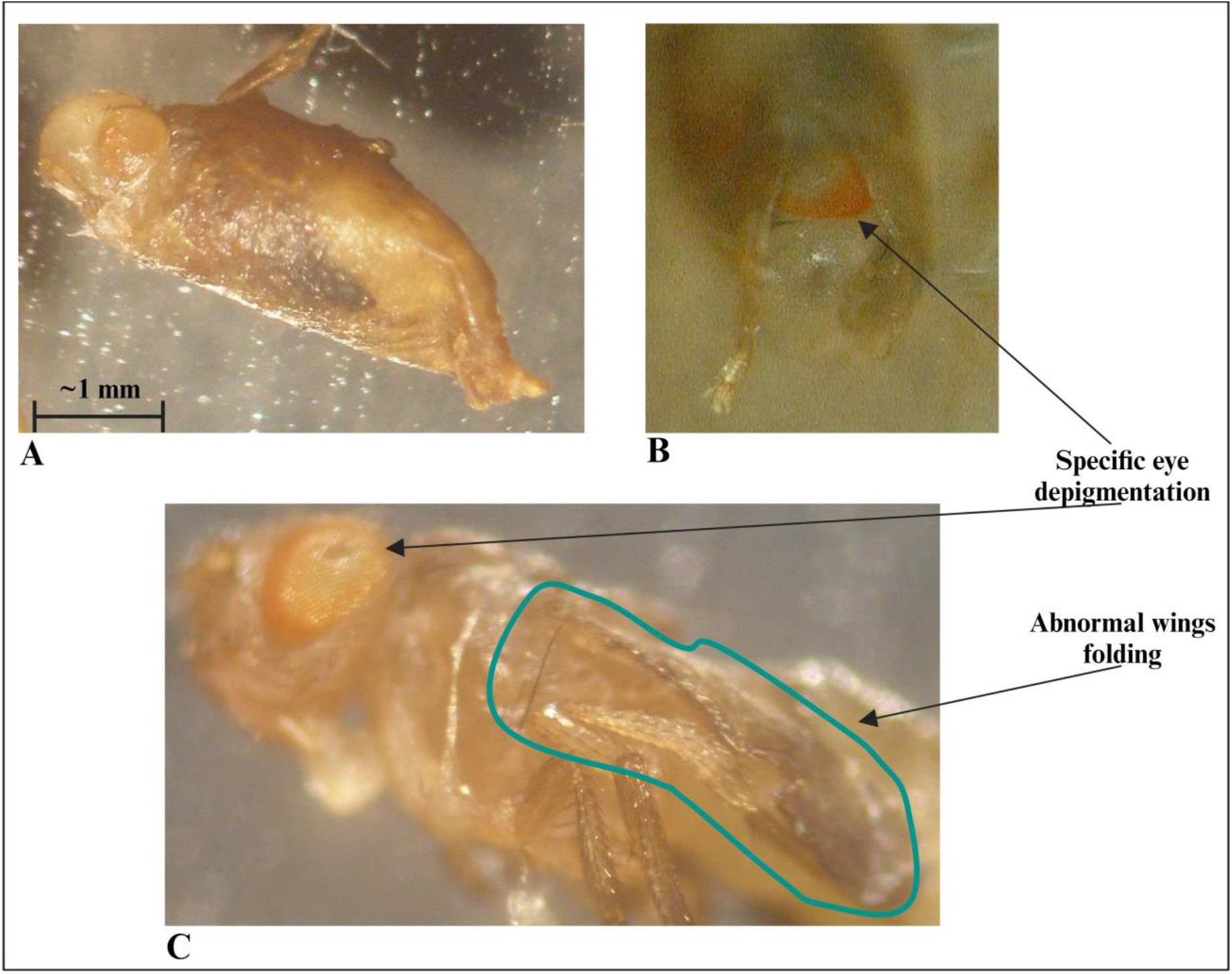
In the figure we highlight the mutant phenotypes characteristic for *CG18135*^*P{lacW}CG18135*^/*CG18135*^*P{lacW}CG18135*^. Most individuals that arrive at the final of the pupal development remain captive within the puparium or only succeed to partially emerge from it (A and B). The imago individuals show a particular eye depigmentation pattern, as well as a persistent abnormal wing folding (C). The horizontal bar shown for Figure 2A indicates the length of one mm.

In addition to these morphological phenotypes, we observed that most adults fail to completely emerge from puparium. Those who are successful are not able to perform locomotion and commonly adopt a stationary position with their legs oriented upwards. Although not able to freely move around, such individuals manifest spasmodic legs movements that apparently manifest in seizure episodes (Supplementary file 2). As previously stated, we did not notice any viable adult older than two days.

### CG18135 gene expression in viable adults

Relative gene expression analysis was based on comparing FC values that were generated with the Livak method. Our data (Figure 3) point no significance for an apparent gene downregulation in *CG18135*^*P{lacW}*^/TM6B mutant females compared to CantonS control females (FC = 1.017) and a significant overexpression of *CG18135* in *CG18135*^*P{lacW}*^/TM6B mutant males versus CantonS control males (FC = 1.850). Moreover, *CG18135* was significantly overexpressed in control males compared to control females (FC = 2.724) with a higher level in mutant males compared to mutant females (FC = 4.927).

**Figure 3.**
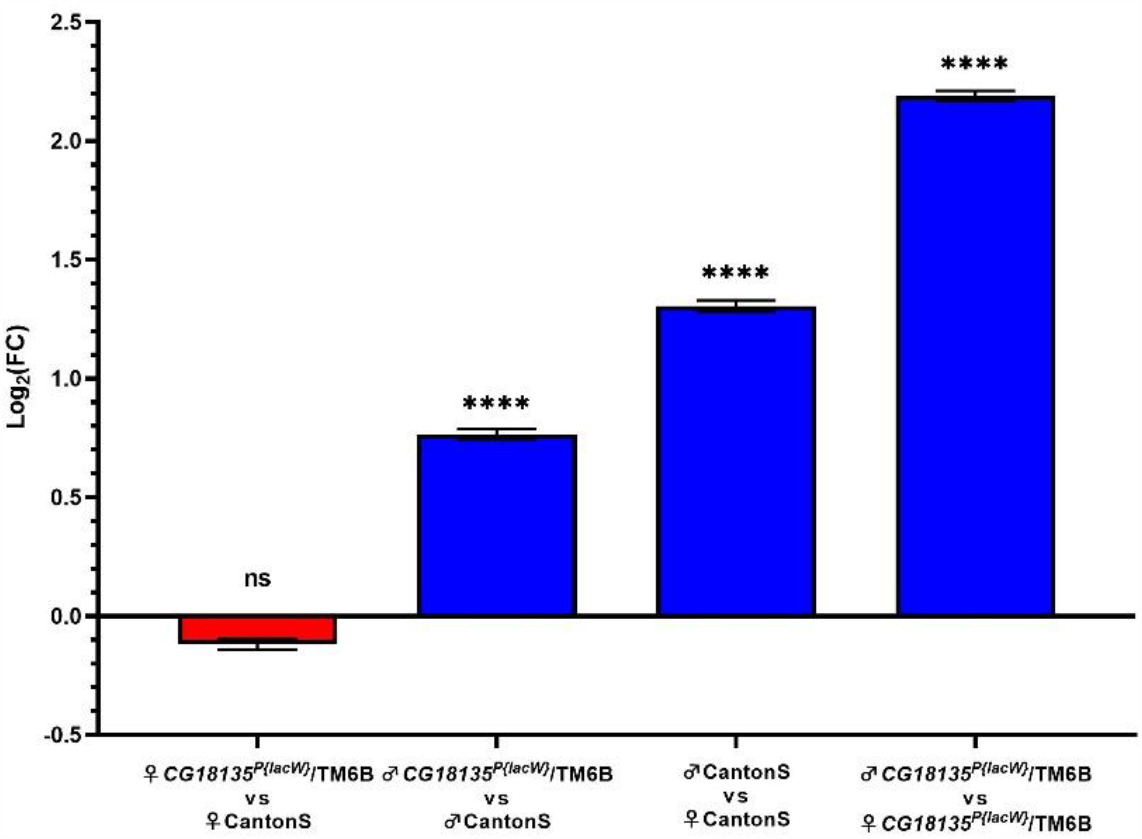
Relative gene expression analysis of *CG18135* between mutant *CG18135*^*P{lacW}*^/TM6B and control CantonS males and females. The bars are in accordance with the log_2_FC values and the ± SEM error bars are shown. Statistical significance was assessed using both Welch two sample t-test and the Exact Wilcoxon rank sum test on 2^-ΔCt^ values corresponding to each analyzed category. A statistically significant difference was considered at p ≤ 0.05 (ns > 0.05, **** < 0.001).

Statistical testing was performed between each experimental group using either the Welch two sample t-test or the Exact Wilcoxon rank sum test with the qDATA application (Ionascu *et al*., 2023). We used these two tests because of the distribution differences of 2^-ΔCt^ values corresponding to the experimental groups. Specifically, while the 2^-ΔCt^ distribution for the heterozygous mutant females was strongly positively skewed, data calculated for the other groups showed a parametric distribution. Moreover, the variance of 2^-ΔCt^ values calculated in the four experimental groups was different within one grade of magnitude, thus suggesting not equal variances. The results are presented in detail in Table 3.

**Table 3.** The results of statistical testing on 2^-ΔCt^ values calculated for mutant *CG18135*^*P{lacW}*^/TM6B and control CantonS groups. Data were obtained with qDATA application.

| Comparative analysis | Welch two sample t-test |  | Exact Wilcoxon rank sum test |  |
| --- | --- | --- | --- | --- |
|  | t statistic | p value | W statistic | p value |
| *CG18135<sup>P{lacW}</sup>/TM6B* ♀ vs CantonS ♀ | -0.826 | 0.415 | 294 | 0.228 |
| *CG18135<sup>P{lacW}</sup>/TM6B* ♂ vs CantonS ♂ | 6.648 | $2.34 \times 10^{-8}$ | 654 | $5.396 \times 10^{-8}$ |
| CantonS ♂ vs CantonS ♀ | 9.138 | $1.4 \times 10^{-10}$ | 715 | $5.218 \times 10^{-13}$ |
| *CG18135<sup>P{lacW}</sup>/TM6B* ♂ vs<br>*CG18135<sup>P{lacW}</sup>/TM6B* ♀ | 15.584 | $3.202 \times 10^{-16}$ | 729 | $1.027 \times 10^{-15}$ |

### The genomic context of CG18135

The nucleotide sequence next to the *P{lacW}*^*CG18135*.*MZ4CM3*^ insertion was aligned against *D. melanogaster* genome (r6.54) using BLAST from FlyBase. The result was visualized in JBrowse with specific tracks selection and a highlight on the CCCTCGCC sequence representing the duplicated octet with the genomic coordinates 3L:18996755..18996762 (Figure 4). This insertion has a type II orientation, meaning that its sense strand is collinear with the minus genomic strand, and it is the only *P{lacW}* hit reported for *CG18135*. Otherwise, only *P, PBac* and *Mimic* derived transposons are reported in FlyBase, such as *P{EPgy2}, P{RS5}, P{XP}, P{EP}*, and *P{SUPor-P}*. Twelve transposon insertions, including *P{lacW}*^*CG18135*.*MZ4CM3*^, span the genomic coordinates 3L:18995536..18996803, which define a genomic region encompassing fragments from the first intron of CG18135-RB and the 5’UTR of CG18135-RA. According to REDfly annotation track, there are four distinct CRMs which overlap or border this genomic region. Out of them, the one denoted REDfly:RFRC:0000014763 (CRM14763) is overlapping the insertion sites of four previously reported *P* derived transposons and of *P{lacW}*^*CG18135*.*MZ4CM3*^. We recovered the CRM14763’s specific sequence and feed it to XSTREAME, a specific tool from the MEME Suite that is able to perform broad motif analysis on sequences with scarce motif sites using generic eukaryote or *D. melanogaster* restricted motif databases. The XTREAME analysis emphasized that CRM14763 harbors specific binding sites for SP1/KLF transcription factors (TFs), MADF-BESS domain transcription regulators and nuclear receptor (ligand-dependent) TFs.

**Figure 4.**
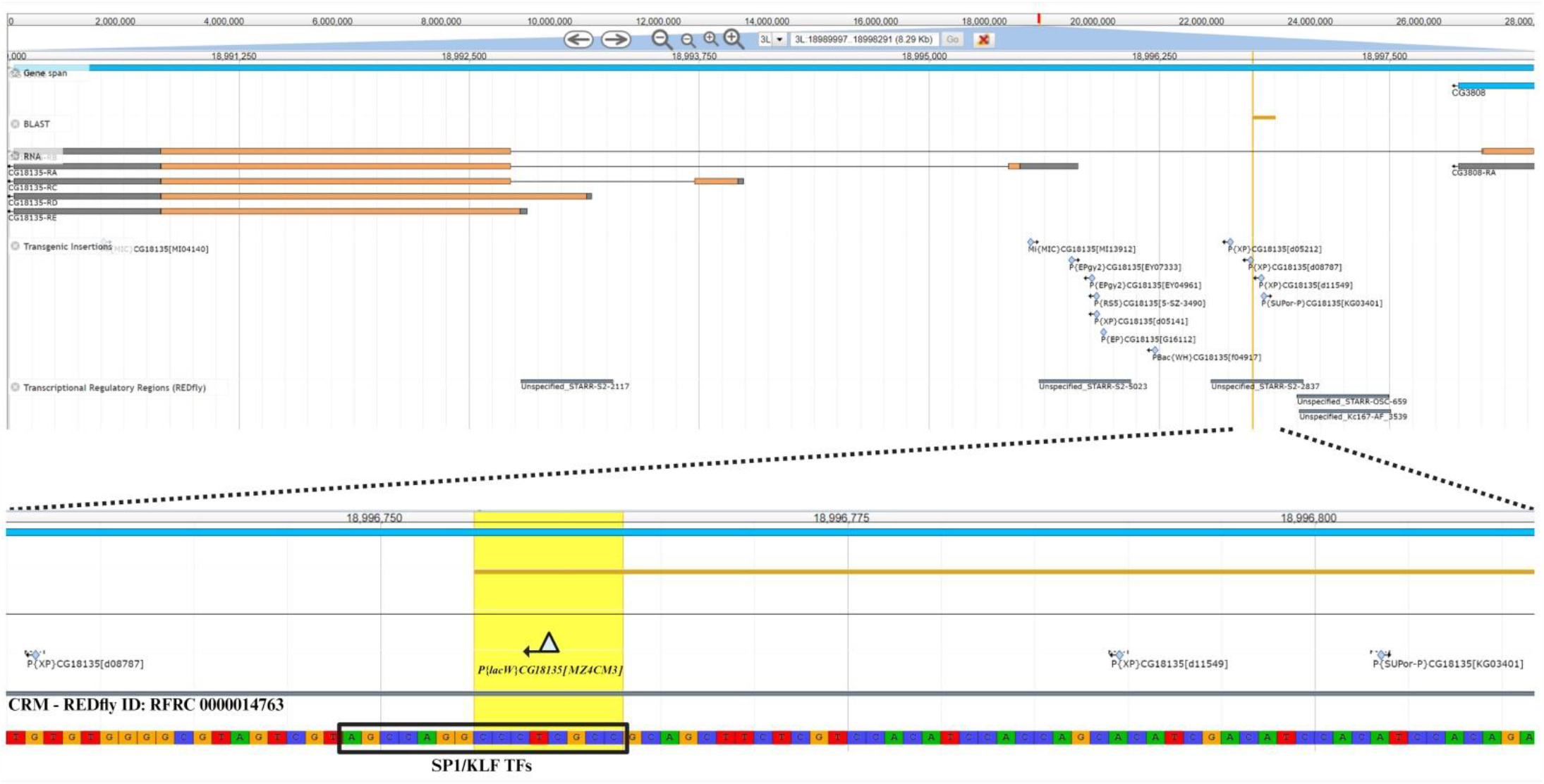
The upper section offers a panoramic view of *CG18135* gene and overlapping genomic features provided by specific JBrowse annotation tracks, such as gene span, RNA, transgenic insertions and transcriptional regulatory regions. In the lower section the *P{lacW}*^*CG18135*.*MZ4CM3*^ insertion, the duplicated octet, as well as its type II orientation are highlighted. The SP1/KLF TFs binding site found inside the CRM with the REDfly ID:0000014763 is indicated as it is located in the innermost vicinity of *P{lacW}*.

Noteworthy, the SP1/KLF TFs’ binding motif that has the AGCCAGG**CCCTCGCC** nucleotide sequence encompasses the sequence duplicated consecutive to *P{lacW}*^*CG18135*.*MZ4CM3*^ insertion (highlighted in bold characters).

## Discussion

In this study we investigated the mutant phenotypes of offspring from *CG18135*^*P{lacW}*^/GFP and *CG18135*^*P{lacW}*^/TM6B lines. We found that, for *CG18135*^*P{lacW}*^/*CG18135*^*P{lacW}*^ mutants, the larval and pupal developmental stages were strongly associated with lethality. To our knowledge, this is the first viability analysis of *CG18135* gene in *D. melanogaster*.

The lack of statistically significant differences between observed and expected lethality of mutant embryos suggests that the *CG18135*^*P{lacW}*^ allele is not associated with embryo lethality. This result could be attributed to the parental delivery of functional CG18135 proteins either in the unfertilized egg (Fisher *et al*., 2012) or via the seminal fluid (Findlay *et al*., 2008), fulfilling the CG18135 protein requirements in early embryonic stages.

Previous proteomics studies identified increased CG18135 protein levels in *D. melanogaster* larvae, pupae and adults, compared to embryos (Brown *et al*., 2014; Casas-Vila *et al*., 2017). Also, high expression of *CG18135* was found in the Bolwig’s organ (Fisher *et al*., 2012) in larvae. The corresponding protein was identified within the plasma membrane of *D. melanogaster* adults’ heads (Aradska *et al*., 2015). Taken together, these data suggest that the CG18135 protein requirement is higher in larvae and, in the absence of wild-type protein production, the maternal or paternal CG18135 in embryos is not sufficient to support normal development, increasing larval lethality.

In addition, we observed that a small percentage of *CG18135*^*P{lacW}*^/*CG18135*^*P{lacW}*^ pupae developed to the adult stage. The CG18135 protein participates in myosin filament binding within muscular tissues, as well as in cell division (Liu *et al*., 2008). Therefore, is expected that the protein requirement increases during pupal development which implies organogenesis and high rates of cellular division. Our previous preliminary data revealed that pupal lethality of CantonS and individuals heterozygous for *CG18135*^*P{lacW}CG18135*^ allele is ≈ 6.7% and ≈ 10.3%, respectively. For *CG18135*^*P{lacW}CG18135*^/*CG18135*^*P{lacW}CG18135*^ mutants, almost all the surviving individuals that get through embryo and larval stages would die inside or attached to the puparium.

These results reveal two major lethality thresholds in the development stages, one in early larval stage (but we do not exclude a certain degree of mortality in the late pre-larval embryonic stage), and a second one in the late stages of pupal development. The latter could be considered an incomplete lethality, since rare imago escapers are active for a maximum of about two days, but beyond that age the lethality induced by the *CG18135*^*P{lacW}CG18135*^/*CG18135*^*P{lacW}CG18135*^ genotype is absolute.

We hypothesize that the *CG18135* gene participates in various biological and molecular processes in *D. melanogaster*, similar to its structural orthologous gene in higher eukaryotes, *GPCPD1*. In humans, *GPCPD1* knockdown malignant cell showed an increased intracellular glycerophosphocholine/phosphocholine (GPC/PC) ratio, a decreased lipid metabolism and a cell migration inhibition (Steward *et al*., 2011). In higher eukaryotes, GPCPD1 facilitates GPC degradation, an essential phospholipidic membrane constituent (Eibl, 1980; Okazaki *et al*., 2010), into glycerol-3-phosphate (G3P) and choline (Steward *et al*., 2012). In eukaryotes, the G3P requirement is obtained from glycolysis (Li *et al*., 2019; Nguyen *et al*., 2019) and, together with choline, it is further used for GPC synthesis via the Kennedy pathway (Veldhuizen *et al*., 1989; Fagone and Jackowski, 2013). These observations suggest that the CG18135 protein may be involved in the regulation of a negative feedback loop between G3P, choline and GPC (Figure 5). Furthermore, choline contributes to *de novo* synthesis of phosphatidylcholine and phosphatidylethanolamine membrane constituents via the Kennedy pathway (Steward *et al*., 2012; Gibellini and Smith, 2010), and participates in Krebs cycle (Steward *et al*., 2012), being an essential component in energy metabolism.

**Figure 5.**
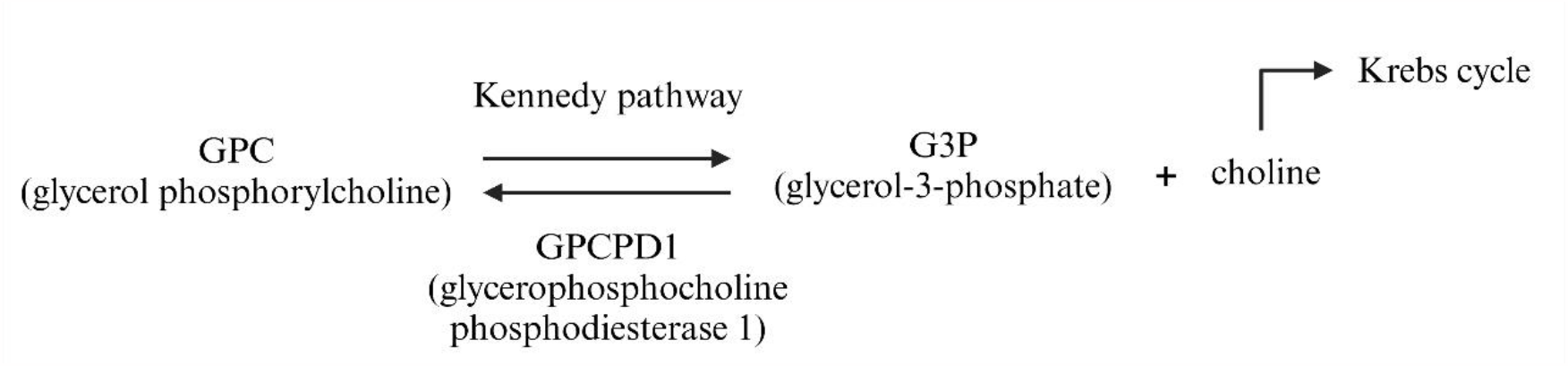
Biochemical pathway of the GPC catabolism mediated by GPCPD1 protein (in humans). It is possible that a similar mechanism is also true for the CG18135 protein in *D. melanogaster*.

The observed phenotypes of homozygous mutant imago escapers are consistent with the proposed involvement of the *CG18135* gene in either muscular activity or energy metabolism.

Participation of *CG18135* in embryo myosin filaments binding has been previously demonstrated (Liu *et al*., 2008). Being localized in the cellular membrane vicinity, *Myo10A* participates in myosin filaments binding during embryogenesis together with several cargo proteins, including DE-cadherin, alpha-tubulin, Katanin-60, milton, atypical protein kinase C (aPKC), Nedd4 and the CG18135 protein. This interaction might be, at least partially, responsible for the unsynchronized movements observed in mutant adult escapers, as the absence of wild-type CG18135 protein could result in muscle contraction abnormalities. In addition, the structural orthologous gene *GPCPD1* has been shown to be associated with muscle development in mice (Okazaki *et al*., 2010, Cikes *et al*., 2022). Okazaki *et al*. (2010) showed that *GPCPD1* gene expression was downregulated in atrophied skeletal muscles, and *GPCPD1* gene knock-out enhances myoblastic differentiation and limits the progression of skeletal muscle atrophy (Okazaki *et al*., 2010). Thus, a lack of sufficient CG18135 protein could increase the lethality rates in larvae or pupae, as observed in our study.

Whilst the correlation between skeletal muscle atrophy and *GPCPD1* expression was previously demonstrated, a recent study suggests that inactivation of *GPCPD1* also results in early muscle ageing due to GPC accumulation (Cikes *et al*., 2022). More specifically, the “aged-like” muscles of *GPCPC1* deficient mice was caused by an abnormal glucose metabolism, and not by muscle development *per se* (Cikes *et al*., 2022). Interestingly, increased GPC levels were found only in muscle (but not in the global lipidome) and did not occur in *GPCPD1* deficiency in other tissues, such as fat, brown fat or liver (Cikes *et al*., 2022). These results suggest that *GPCPD1* is associated with glucose metabolism within the muscle tissue.

Besides from the spasmodic leg movement of imago escapers, we observed a general lack of activity, which could be caused by either an abnormal energy metabolism or by a decreased response to environmental stimuli. We would rather overlook the latter, as *CG18135*^*P{lacW}*^/GFP is derived from a line that did not show any phenotypes that might suggest abnormal response to stimuli. Considering the previously discussed functions of *GPCPD1* in muscle contractions and energy metabolism, the sedentary phenotype might be caused by a lack of global energy availability in the imago escapers, or by a lack of coordinated muscle contractions which require ATP. These phenotypes were evident in imago individuals, but we did not notice such phenotypes in larvae. Although more investigations are required, preliminary data might suggest that CG18135 protein necessity is higher in the later stages of development. Further experiments might focus on the expression of *CG18135* gene in mutant larvae or pupae, but also on the GPC/PC ratio in mutants compared to wild-type.

Considering the high lethality of *CG18135*^*P{lacW}CG18135*^*/CG18135*^*P{lacW}CG18135*^ mutant flies prior to adult stage, we decided to evaluate the *CG18135* gene expression in adult flies, both in male versus female groups and in mutant versus control groups. Following the quantification of *CG18135* gene expression in different *D. melanogaster* adult groups, we found that *CG18135*^*P{lacW}CG18135*^/TM6B mutant females and CantonS control females displayed a similar expression level, even if the mutant females had only one functional allele of *CG18135*.

Interestingly, we found that *CG18135* was significantly overexpressed in mutant males compared to control males. It is plausible that in the absence of one functional allele there might be an increase in the GPC/PC ratio (Steward *et al*., 2012) which may result in greater CG18135 protein requirement in order to reduce the GPC/PC ratio, and an upregulation of *CG18135*. Nonetheless, because biochemical pathways involving phospholipids are complex, and because we did not observe similar expression profiles in mutant females compared to control females, it is not possible to infer a definitive conclusion.

In males, the CG18135 protein is present in the seminal fluid of *D. melanogaster* individuals at mating (Findlay *et al*., 2008), which could partially explain the overexpression of *CG18135* gene in mutant or wild-type males compared to females. Additionally, the reproductive system of male flies is associated with intense cellular division and cellular differentiation during spermatogenesis (Fabian and Brill, 2012). These processes are dependent on glycerophospholipids availability and may influence the expression of *CG18135*.

The observed *CG18135* overexpression in males compared to female adult flies is consistent with previous transcriptomics studies (Brown *et al*., 2014; Casas-Vila *et al*., 2017). The *CG18135* overexpression in mutant males versus mutant females (FC = 4.927) could be caused by a combination of factors that drive overexpression in mutant males compared to control males (FC = 1.85) and in control males compared to control females (FC = 2.724).

We intend to evaluate targeted qRT-PCR gene expression of putative interactors of *CG18135*. Candidate genes include *Myo10A* and its neighboring orthologue *CG11619* gene, as well as specific genes acting downstream or upstream of the glycerophospholipid metabolism, such as *choline acetyltransferase* (*ChAT*) and *protein kinase c-AMP dependent regulatory subunit 1* (*Pka-R1*), respectively.

Genomic context analysis showed that the *P{lacW}* insertion in *CG18135* resides adjacently to the binding site of TF *cabut* (*cbt*), which acts upstream of *decapentaplegic* (*dpp*) in their respective biochemical pathway. As a consequence, it is possible that the binding site is not efficiently accessed by the TF. Therefore, a series of downstream interactions are influenced, including dorsal closure in embryos, wing disc morphogenesis, muscle degeneration, and energy metabolism. The abnormal *cbt* activation in our mutants might partially explain some of the observed phenotypes. For example, *cbt* overexpression in embryos was found only in later stages embryo (but not in early ones) in the neurons and glia cells of the central nervous system (Belacortu *et al*., 2011). Although we did not observe significant embryo lethality, *cbt* interaction with *CG18135* might be supported by the identification of *CG18135* in the phospholipid membranes in adult flies heads (Aradska *et al*., 2015). Wing disc morphogenesis involvement of *cbt* (Benjarano *et al*., 2008; Rodriguez, 2011) might be correlated with the observed abnormal wing phenotype in mutant imago escapers. Moreover, *cbt* has been recently considered to influence the mechanism of flight and jump muscles control and also to regulate the number of tergal depressor of the trochanter (TDT) jump muscles (Giedd, 2018). Along with the findings of a *cbt* link to energy metabolism via a regulatory link between sugar sensing and the circadian clock (Bartok *et al*., 2015), *cbt* interaction with the *CG18135* transcription site might further aggravate the sedentary behavior or sporadic movements of mutant adult escapers.

In humans, the homologues of *cbt* are the TFs *Krüppel-like factor 10* (*KLF10*) and *KLF11* (Bartok *et al*., 2015). Both TFs are part of a TFs family that regulate CG-rich promoters (Lomberk *et al*., 2012) and influence the energy metabolism. Whilst *KLF10* is overexpressed in response to high sugar levels (Iizuka *et al*., 2011), *KLF11* is upregulated during starvation (Zhang *et al*., 2011) and it has been suggested that the former is a true *cbt* functional orthologue (Bartok *et al*., 2015). These TFs, also known as *TGFβ-inducible early gene 1* (*TIEG1*) and *TIEG2*, respectively, are involved in *TGF-β* mediated cell proliferation, differentiation and apoptosis (Lomberk and Urrutia, 2005). What is more, *KLF* TFs family has been shown to downregulate the activity of *Sp1 transcription factor* (*SP1*), a ubiquitous TF that regulates hundreds of genes (Lomberk and Urrutia, 2005). In our view, an abnormal binding of *cbt* to the *CG18135* site might exert even a more profound effect over the *CG18135* overall transcription regulation, since various CRMs that could share functional roles are forcibly separated. This evidence supports the importance of normal *cbt* activity in the cell and highlights the relevance of our *D. melanogaster* mutant lines.

## Conclusions

In this study we characterized for the first time mutant phenotypes produced by the *CG18135*^*P{lacW}CG18135*^ insertional allele in *D. melanogaster*. We performed several viability assessments allowing us to establish that the early larval and late pupal developmental stages are critical for lethality onset. Rare imago escapers that live a very short life exhibit very specific phenotypes affecting the eye and wing morphology, as well as their locomotor proficiency. These severe phenotypes are probably a consequence of the regulatory perturbations that are likely to be caused by the insertion of *P{lacW}*^*CG18135*.*MZ4CM3*^ transposon inside a group of CRMs containing binding sites recognized by wing and energy metabolism specific, as well as ubiquitous TFs. Whilst heterozygous flies are apparently normally developing, the gene expression of *CG18135* is significantly different in male compared to female flies and in mutant compared to control males. Our findings support the essential role of *CG18135* gene in *D. melanogaster* and define a novel experimental platform for modelling the interactions of *GPCPD1* structural and functional orthologue gene in the higher eukaryotes.

## Acknowledgments

This work was funded by the Academy of Romanian Scientists as part of the research grant number AOSR-TEAMS II, 2023-2024 Edition, No. 1 - Biological Sciences Domain.

